# Chemogenetic silencing of neurons in the mouse anterior cingulate area modulates neuronal activity and functional connectivity

**DOI:** 10.1101/686725

**Authors:** Lore M. Peeters, Rukun Hinz, Jan R. Detrez, Stephan Missault, Winnok H. De Vos, Marleen Verhoye, Annemie Van der Linden, Georgios A. Keliris

## Abstract

The anterior cingulate area (ACA) is an integral part of the prefrontal cortex in mice and has been implicated in several cognitive functions. Previous anatomical and functional imaging studies demonstrated that the ACA is highly interconnected with numerous brain regions acting as a hub region in functional networks. However, the importance of the ACA in regulating functional network activity and connectivity remains to be elucidated. Recently developed neuromodulatory techniques, such as Designer Receptors Exclusively Activated by Designer Drugs (DREADDs) allow for precise control of neuronal activity. In this study, we used an inhibitory kappa-opioid receptor DREADDs (KORD) to temporally inhibit neuronal firing in the right ACA of mice and assessed functional network activity and connectivity using non-invasive functional MRI. We demonstrated that KORD-induced inhibition of the right ACA induced blood oxygenation-level dependent (BOLD) signal decreases and increases in connected brain regions throughout of hemispheres. Furthermore, these modulations in neuronal activity were associated with decreased intra- and interhemispheric functional connectivity. These results demonstrate that the combination of the DREADD technology and non-invasive functional imaging methods is a valuable tool for unraveling the underlying mechanisms of network function and dysfunction.

## 1. Introduction

The anterior cingulate area (ACA) is an integral part of the prefrontal cortex and has been implicated in several cognitive functions including attention [1-3], remote memory [4, 5], motion planning and execution [6] and processing of pain [7]. Accumulating evidence suggests that the cytoarchitecture and functional role of the ACA is similar across different species including humans, non-human primates and rodents [8-11]. Anatomical and functional imaging studies identified the ACA as a central hub region that is highly interconnected with numerous brain regions and involved in multiple functional networks [12-14]. ACA hypoconnectivity, as measured in functional MRI studies, is associated with network dysfunctions implicated in multiple psychiatric disorders, such as attention-deficit hyperactivity disorder [15], schizophrenia [16, 17], bipolar disorder [18, 19], depression and anxiety [20]. Furthermore, abnormal functional connectivity (FC) of the ACA has also been shown to occur following traumatic brain injury [21, 22]. This implicates that the ACA is involved in a wide range of neurological disorders. Nevertheless, the direct role of neuronal populations in the ACA in maintaining neuronal network function has not yet been examined to date.

During the last decade, researchers have focused on the development of new neuromodulatory tools, such as optogenetics and chemogenetics, which allow controlling the activity of specifically targeted neuronal populations with spatiotemporal specificity. Current chemogenetic tools include Designer Receptors Exclusively Activated by Designer Drugs (DREADDs), which are engineered receptors for targeted enhancement or silencing of neurons upon binding of an otherwise inert ligand. The combination of chemogenetics and non-invasive neuroimaging methods, such as functional magnetic resonance imaging (fMRI), allows direct evaluation of DREADD-induced changes in neuronal firing on large-scale neuronal activity and FC. As such, Giorgi et al. were able to causally link the chemogenetic activation of serotonergic neurons with neuronal activity increases in cortical and subcortical brain areas, measured by relative cerebral blood volume increases [23]. Furthermore, other studies also used blood-oxygenation-level-dependent (BOLD) fMRI to identify the effect of chemogenetic activation of mesolimbic and mesocortical pathways [24] or chemogenetic inhibition of the amygdala [25] on whole-brain network integrity.

Here, we combined inhibitory kappa opioid-receptor DREADDs (KORD) in the mouse ACA and non-invasive fMRI in order to evaluate the link between inhibition of the ACA and whole-brain network activity and connectivity. We hypothesized that acute inhibition of the ACA leads to neuronal activity and FC alterations in structurally and functionally connected brain regions. Pharmacological MRI (phMRI) was used to assess changes in BOLD activation levels after injection of the chemogenetic agent, Salvinorin B (SalB), an inert ligand of the KORD. Changes in neuronal activity were evaluated in the target region as well as in connected brain areas. Resting state fMRI (rsfMRI) was used to identify brain wide FC alterations that may reflect neuronal network reorganizations and disruptions similar to disease states.

## 2. Methods

### 2.1 Animals and ethics statement

Healthy male C57BL/6J mice (N = 18; Janvier, France) of (34 ± 1) g were used in the study. Animals were group housed under controlled humidity (40%) and temperature (20 – 24°C) conditions, with a 12h light/dark cycle. Standard food and water were provided ad libitum. All procedures were in accordance with the guidelines approved by the European Ethics Committee (decree 2010/63/EU) and were approved by the Committee on Animal Care and Use at the University of Antwerp, Belgium (approval number: 2016-49).

### 2.2 Intracerebral viral vector injections

Mice were subjected to a stereotactic surgery targeting the right ACA (AP +0.86 mm, ML +0.50 mm, DV −0.75 mm). Mice were randomly divided into two groups: a KORD-expressing group (N=12) and a sham group (N=6). Viral vector injections were performed as follows: mice were anesthetized using 2% isoflurane (Isoflo^®^, Abbott Laboratories Ltd., USA) and received a subcutaneous (s.c.) injection of xylocaine (Lidocaine hydrochloride, Astra Zeneca) for local analgesia. The following viral vectors were injected: 1 µL of 3.3 x 10^12^ vector genomes (vg)/mL of AAV8-CaMKII-HA-KORD-IRES-mCitrine (Vector Core, University of North Carolina, USA) in the KORD-expressing group, and 1 µL of 5.6 x 10^12^ vg/mL of AAV8-CaMKIIa-EGFP (Vector Core, University of North Carolina, USA) in the sham group. The virus was injected at a rate of 18.3 nL/min with a nano-injector (Nanoject II, Drummond). The MRI experiments started at least three weeks after viral vector injection to allow optimal expression levels [26].

### 2.3 In vivo MRI procedures

Initially, mice were anesthetized with 2% isoflurane in a mixture of 70% N_2_ and 30% O_2_. During the imaging procedures, an optimized anesthesia protocol was used [27]. Briefly, a s.c. bolus injection of 0.05 mg/kg medetomidine (Domitor^®^, Pfizer, Germany) was administered, and after 10 min a s.c. infusion of 0.1 mg/kg/h medetomidine was started. Meanwhile, the isoflurane level was gradually decreased to 0.3%. Functional imaging scans were acquired starting from forty minutes post-bolus injection. The animals’ physiology was closely monitored throughout the acquisition of the MRI scans. The breathing rate was recorded using a pressure-sensitive pad (sampling rate 225 Hz; MR-compatible Small Animal Monitoring and Gating system, SA Instruments, Inc., USA) and the body temperature was maintained at (37 ± 0.5) °C using a feedback controlled warm air circuitry (MR-compatible Small Animal Heating System, SA Instruments, Inc., USA). In addition, the level of blood oxygenation was monitored using a pulse oximeter (MR-compatible Small Animal Monitoring and Gating System, SA Instruments, Inc., USA). After the imaging procedures, the medetomidine anesthesia was reversed by injecting 0.1 mg/kg atipamezole (Antisedan^®^, Pfizer, Germany).

All MRI scans were acquired on a 9.4T Biospec MRI system (Bruker BioSpin, Germany) with Paravision 6.0.1 software. The images were acquired using a standard Bruker crosscoil set-up with a quadrature volume coil and a quadrature surface coil designed for mice. First, three orthogonal multi-slice Turbo RARE T_2_-weighted images were acquired to enable uniform slice-positioning (TR: 2000 ms, TE: 33 ms, matrix: [256×256], FOV: (20×20) mm^2^, in-plane resolution: (0.078×0.078) mm^2^, 16 coronal slices of 0.4 mm). Then, inhomogeneity of the magnetic field in an ellipsoidal volume of interest within the brain was corrected by local shimming. The phMRI scans and rsfMRI scans were acquired during separate scan sessions.

### 2.4 Assessing KORD-induced neuronal activity changes

#### 2.4.1 MRI data Acquisition

PhMRI data were acquired using a T_2_ ^*^weighted echo planar imaging (EPI) sequence (TR: 15000 ms, TE: 20 ms, matrix: [64×64], FOV: (20×20) mm^2^, two segments, in-plane resolution: (0.312×0.312) mm^2^, 16 coronal slices of 0.4 mm covering cerebrum excluding olfactory bulb). Baseline scans were acquired for 10 min (20 volumes) after which SalB (3 mg/kg (sham and KORD group) or 6 mg/kg (KORD group); Toronto Research Chemicals, Canada; dissolved in DMSO, as previously described in [26]) was administered through a s.c. catheter. Following the acquisition of the ten minutes baseline scan, the scan was continued until fifty minutes (100 volumes) after the SalB injection. The KORD group was scanned with different SalB concentrations (3 mg/kg and 6 mg/kg, separate scan sessions) to assess concentration effects. A total of three scans, two scans from the sham group and one scan from the 3 mg/kg SalB KORD group, were excluded from the phMRI analysis due to signal instabilities. Final group sizes for the phMRI scans were: sham group: N=4 and KORD-expressing group: 3mg/kg SalB N=11, 6 mg/kg SalB N=12.

#### 2.4.2 MRI data pre-processing

Data pre-processing was performed using SPM12 software in MATLAB 2014a. Realignment and unwarping were performed using a least-squares approach and a 6-parameter rigid body spatial transformation. Second, all scans were normalized to the study-specific mean EPI template using a global 12-parameter affine transformation, followed by a non-linear transformation. Then, in-plane smoothing was performed on masked images with a Gaussian kernel with full width at half maximum of twice the voxel size.

#### 2.4.3 MRI data analysis

The scans of the sham group were used to assess potential non-specific effects of SalB and its solvent DMSO on the BOLD signal. Previous studies reported that SalB is an inactive metabolite of the KOR-selective agonist Salvinorin A [26, 28]. However, DMSO has been shown to be able to modulate central nervous system function [29]. To estimate this effect, we used the sham group that didn’t express designer receptors that could be affected by SalB/DMSO. Specifically, the BOLD signal from each voxel was converted to percent signal change using the pre-injection baseline volumes (volumes 1-20) as a reference. Then, the percent BOLD signal change across all sham animals were voxel-wise averaged to create an average sham percent BOLD signal change scan. The resulting average percent BOLD signal change timeseries showed a small linear signal drift before the injection (commonly observed drift in MRI imaging), while after the injection an exponential-like decline indicated unspecific effects of SalB/DMSO. These effects were fitted for each voxel by a linear fit with parameters a, b (volumes 5-20, given by equation (1)) and a double exponential fit with parameters c, d, f and g (volumes 25-120, given by equation (2)), which were connected at their crossing point to determine the final fitted curves (see Supplementary figure 1):

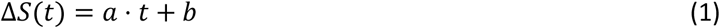

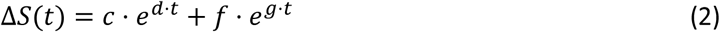

To identify the specific effects of the SalB injection in the KORD groups we calculated the differences between the timeseries (percent signal change) of the KORD-expressing animals and the averaged sham fits. To be certain that our fitting procedures closely reflected the BOLD activity in the sham group, we only considered voxels with very high goodness of fit (R^2^ ≥ 0.85, Supplementary figure 1). Then, the area under the curve (AUC) for the period starting at 10 min after SalB injection until 50 min post-injection (volumes 40-120) was calculated in the KORD groups and used in further statistical analyses. For each SalB concentration, mean statistical maps (one sample Student’s T-test) were computed to test whether the AUC values were statistically different from zero. In addition, a Paired Student’s T-test was performed to compare the effects of the two SalB concentrations in the KORD scan sessions. Next, regions of interest (ROIs) were selected based on the Franklin and Paxinos mouse brain atlas (third edition) in various brain regions, including: anterior cingulate area (ACA), retrosplenial cortex (RSP), insular cortex (Ins), visual cortex (VIS), somatosensory cortex (SS), auditory cortex (AUD) / temporal association cortex (TEA), amygdalar nuclei (AN) and hippocampus (HIP) (Supplementary Figure 2).

### 2.5 Investigating KORD-induced functional connectivity changes

#### 2.5.1 MRI data acquisition

Resting state scans were acquired using a T_2_^*^ weighted single shot EPI sequence (TR: 2000 ms, TE: 16 ms, matrix: [128×64], FOV: (20×20) mm^2^, in-plane resolution: (0.156×0.312) mm^2^, 16 coronal slices of 0.4 mm covering cerebrum excluding olfactory bulb). Two consecutive five-minute rsfMRI scans (150 volumes each) were acquired starting ten minutes after the injection of either vehicle (DMSO) or SalB (3mg/kg). The vehicle and SalB were used in MRI sessions. For the analysis, we selected the first of the two five-minute scans unless excessive motion was observed, in which case the second scan was used (N=2). The rsfMRI scans were acquired only in the KORD-expressing group.

#### 2.5.2 MRI data pre-processing

The rsfMRI scans were pre-processed using SPM12 software in MATLAB2014a. Realignment was performed using a least-squares approach and a 6-parameter rigid body spatial transformation. Further pre-processing steps, consisting of normalization and smoothing, are analogous to the phMRI pre-processing steps described above. The REST toolbox (REST 1.8, http://resting-fmri.sourceforge.net) was used to filter the rsfMRI data. The band-pass filter was set between 0.01 and 0.1Hz to retain the low frequency fluctuations of the BOLD signals.

#### 2.5.3 MRI data analysis

Seed-based analyses were performed to assess FC alterations between brain regions that are known to anatomically or functionally connect to the ACA. As such, seeds, consisting of 4 voxels, were drawn in the ACA, RSP, Ins, VIS, SS, AUD/TEA, HIP and thalamus (Thal) for both hemispheres separately (Supplementary Figure 3). The REST toolbox was used to obtain the mean BOLD signal time course of a seed region. Then, this temporal signal was used in a general linear model analysis to compare it to the timecourse of all other voxels in the brain. This resulted in a FC map consisting of voxels that significantly correlate with the temporal signal of the seed region. Group mean FC maps were calculated for each seed region (One sample Student’s T-test). Masks were created consisting of the significant clusters of the mean group FC maps of the vehicle scans and SalB scans (uncorrected p≤0.001, minimal cluster size = 10 voxels). The masks of both scans were summed and these total masks were used in further analyses. Mean T-values were extracted from the total masks. A Paired Student’s T-test was used to compare the mean T-values between both conditions. To assess interhemispheric FC, ROI time courses were extracted and Pearson’s correlation coefficients were calculated for each pair of ROIs. Then, the correlation values were z-transformed and differences between the two conditions were assessed using a paired Student’s T-test.

### 2.6 Whole-brain microscopy

#### 2.6.1 Clearing and image acquisition

After the in vivo imaging procedures, brain samples were collected to evaluate KORD expression. Mice were deeply anesthetized by an intraperitoneal injection with pentobarbital (Dolethal^®^, Vetoquinol, Belgium). Cardiac perfusion, brain tissue preparation and clearing were done using the uDISCO protocol [30]. Cleared mouse brains were recorded on an Ultramicroscope II (Lavision Biotec GmbH), equipped with an Olympus MVPLAPO 2X (NA 0.50) objective lens and DBE-corrected LV OM DCC20 dipping cap. Images were acquired with a Neo sCMOS camera (Andor) at 1.6X magnification and a 10µm z-step resulting in a voxel size of 2 µm x 2 µm x 10 µm. Images acquired with left and right light sheet illumination were merged with a linear blending algorithm. A 488nm (for EGFP and mCitrine) and 561nm laser (for autofluorescence) with a 525/50nm and 620/60nm emission filters were used.

#### 2.6.2 Image analysis

Regional analysis of the EGFP and mCitrine expression was done as previously described [31]. Briefly, the autofluorescence channel was aligned to a 3D autofluorescence reference brain atlas (Allen Brain atlas) using Elastix, and the resulting transformation vector was used for regional analysis of the fluorescent protein signal [32]. To segment the fluorescent protein signal, a ratio image was generated by dividing the signal channel by the autofluorescent channel in FIJI [33]. Edges were subsequently enhanced by applying a Laplace filter (radius 75 µm), and binarized using a user-defined intensity threshold. Spurious signals (e.g. at the contours of the brain, and edges within the tissue) were manually removed. Finally, the total number of detected voxels for a given brain region volume was calculated and expressed relative to the brain region size (Supplementary Figure 4).

## 3. Results

Post-mortem whole-brain imaging after brain clearing (uDISCO) demonstrated that the targeted injection site overlaps with the ACA region of the right hemisphere in both the sham group and the KORD-expressing group (Supplementary Figure 4). Regional quantification of the mCitrine fluorescence showed that significant signal from the KORD expression could only be observed in the target region. However, similar injections of the EGFP control virus elicited stronger fluorescence that in addition to the target region could also be observed in projecting regions, including motor and somatosensory regions, retrosplenial cortex, insular cortex, thalamic nuclei and caudate putamen, with spreading to the contralateral hemisphere.

### 3.1 Large scale network modulations upon KORD-induced neuronal inhibition in the right anterior cingulate area

Pharmacological MRI was performed to assess in which brain regions the neural activity was modified upon chemogenetic inhibition of the neurons in the right ACA. The mean statistical difference maps from the comparison of both SalB concentrations in the KORD-expressing mice showed no significant clusters (Paired Student’s T-test, p≤0.005 uncorrected, minimal cluster size = 10 voxels). Therefore, the scans of both conditions were grouped to evaluate the effect of SalB on the BOLD signal. Upon SalB injection, the mean statistical AUC maps showed both significant BOLD signal decreases as well as increases (One sample Student’s T-test, p≤0.005, minimal cluster size = 10 voxels, Figure 1). Decreased BOLD signals were observed as expected in the target region ACA, and in addition in Ins, left SS, left AUD/TEA and right VIS, while, increased BOLD signals were observed in the right RSP, right SS, right HIP and AN. The mean AUC values ± SEM from all voxels in the brain regions demonstrating significant decreases or increases (One sample Student’s T-test) are presented in Table 1.

**Figure 1.**
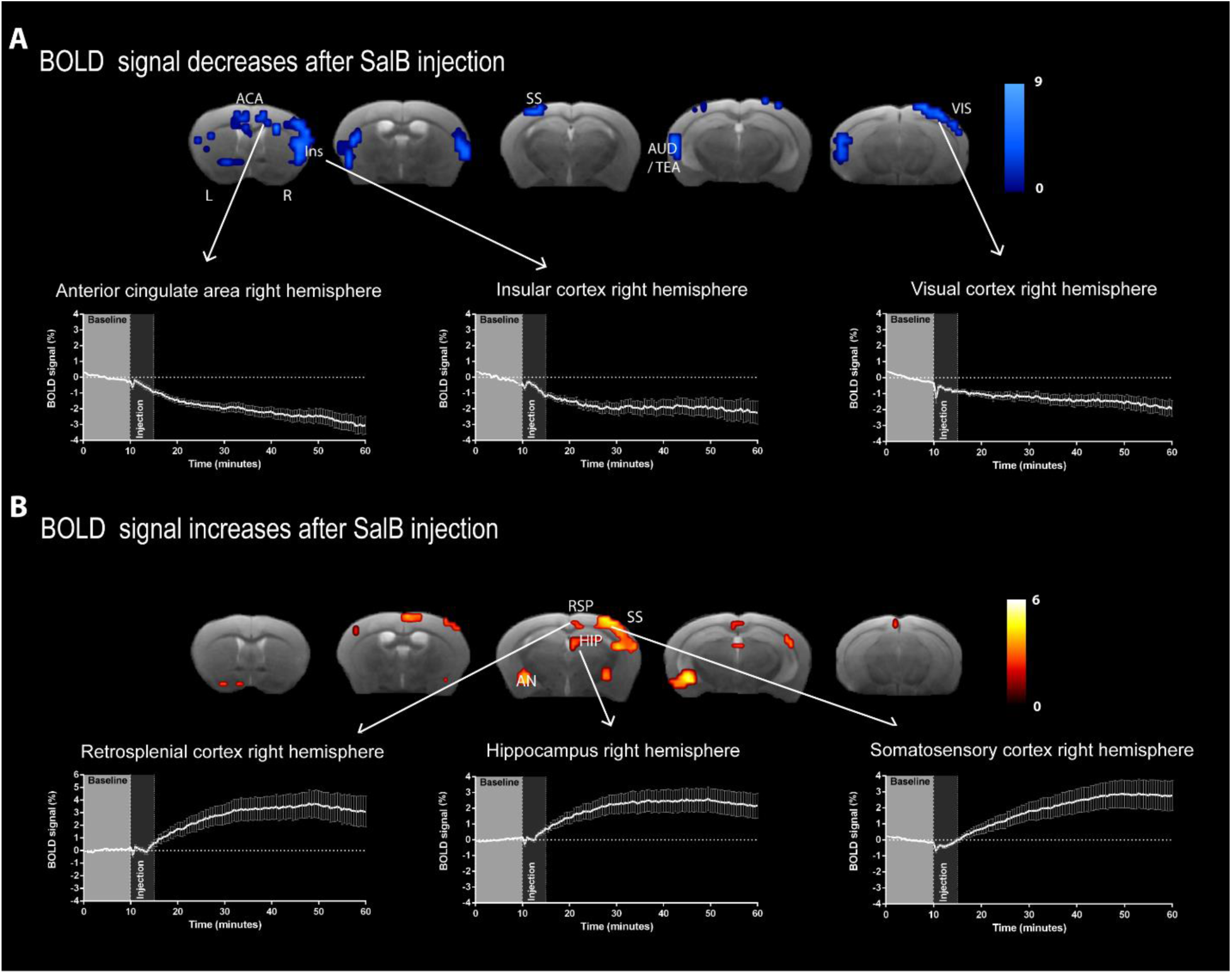
Neural activity alterations upon neuronal inhibition in right ACA. Mean statistical AUC maps in the KORD-expressing group are shown. A) Mean statistical AUC maps of the significantly decreased BOLD signals after SalB injection. Graphs show the BOLD signal timeseries of all significant voxels in the right ACA, right Ins and right VIS. B) Mean statistical AUC maps of significantly increased BOLD signals after SalB injection. Graphs show the percent BOLD signal change timeseries of all significant voxels in the right RSP, right HIP and right SS. One sample T-test, p≤0.005, minimal cluster size = 10 voxels. Abbreviations: anterior cingulate area (ACA), insular cortex (Ins), auditory cortex (AUD), temporal association cortex (TEA), visual cortex (VIS), retrosplenial cortex (RSP), amygdalar nuclei (AN), somatosensory cortex (SS), hippocampus (HIP).

**Table 1.** Quantification of the area under the curve (AUC). Table shows the mean AUC ± SEM from the extracted timecourses of the significant voxels of different ROIs, including anterior cingulate area (ACA), insular cortex (Ins), somatosensory cortex (SS), auditory cortex (AUD) / temporal association cortex (TEA), hippocampus (HIP), retrosplenial cortex (RSP), visual cortex (VIS), amygdalar nucleus (AN).

| Hemisphere | Brain region | Mean AUC ± SEM |
| --- | --- | --- |
| Left hemisphere | Anterior cingulate area (ACA) | -1,3 ± 0,3 |
| | Insular cortex (Ins) | -1,5 ± 0,4 |
| | Somatosensory cortex (SS) | -1,5 ± 0,3 |
| | Auditory cortex / temporal association cortex (AUD/TEA) | -2,1 ± 0,5 |
| | Amygdalar nucleus (AN) | 3,0 ± 0,8 |
| Right hemisphere | Anterior cingulate area (ACA) | -1,2 ± 0,4 |
| | Insular cortex (Ins) | -2,4 ± 0,5 |
| | Somatosensory cortex (SS) | 2,7 ± 0,7 |
| | Hippocampus (HIP) | 2,0 ± 0,7 |
| | Retrosplenial cortex (RSP) | 2,5 ± 0,8 |
| | Visual cortex (VIS) | -1,4 ± 0,3 |
| | Amygdalar nucleus (AN) | 3,4 ± 1,0 |

### 3.2 KORD-induced functional connectivity reductions

To evaluate how functional connectivity between different brain regions in changing during KORD-induced inactivation of the ACA, we analyzed the rsfMRI scans that were collected ten minutes after the injection of SalB. Seed-based analyses revealed that inhibition of the neurons in the right ACA significantly decreased FC of the right and left ACA, right and left RSP, right SS, right and left VIS, right AUD, left HIP and right and left Thal (Paired Student’s T-test, Figure 2). Furthermore, interhemispheric FC was assessed between homologous ROIs in both hemispheres. Significantly decreased interhemispheric FC was measured for the RSP, SS and VIS (Paired Student’s T-test, Figure 3).

**Figure 2.**
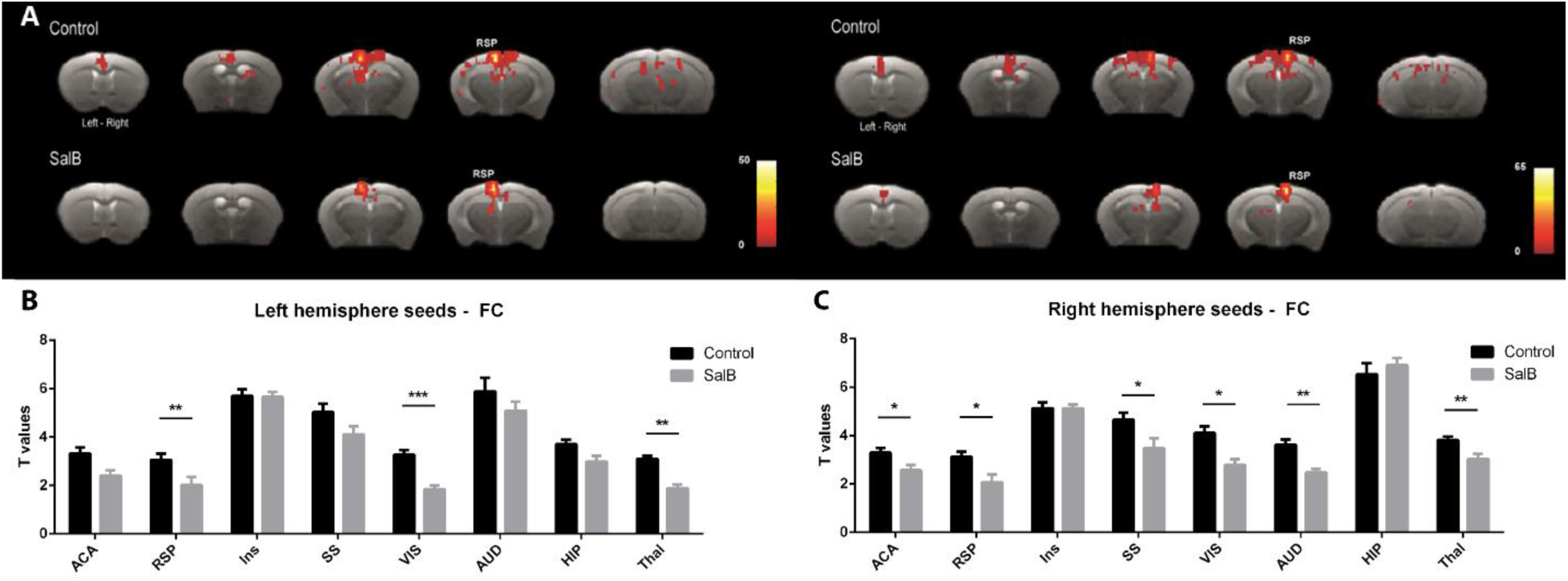
Functional connectivity alterations due to inhibition of the right ACA. A) Statistical FC maps of the RSP as seed region after vehicle injection (control) and SalB injection (One sample T-test, p≤0.001 uncorrected, minimal cluster size 10 voxels). B + C) Bar graphs show the quantification of the mean T-values ± SEM, i.e. FC strength, for all seed regions (anterior cingulate area (ACA), retrosplenial cortex (RSP), insular cortex (Ins), visual cortex (VIS), somatosensory cortex (SS), auditory cortex (AUD), hippocampus (HIP) and thalamus (Thal); Paired Student’s T-test). * p≤0.05, ** p≤0.01, *** p≤0.001.

**Figure 3.**
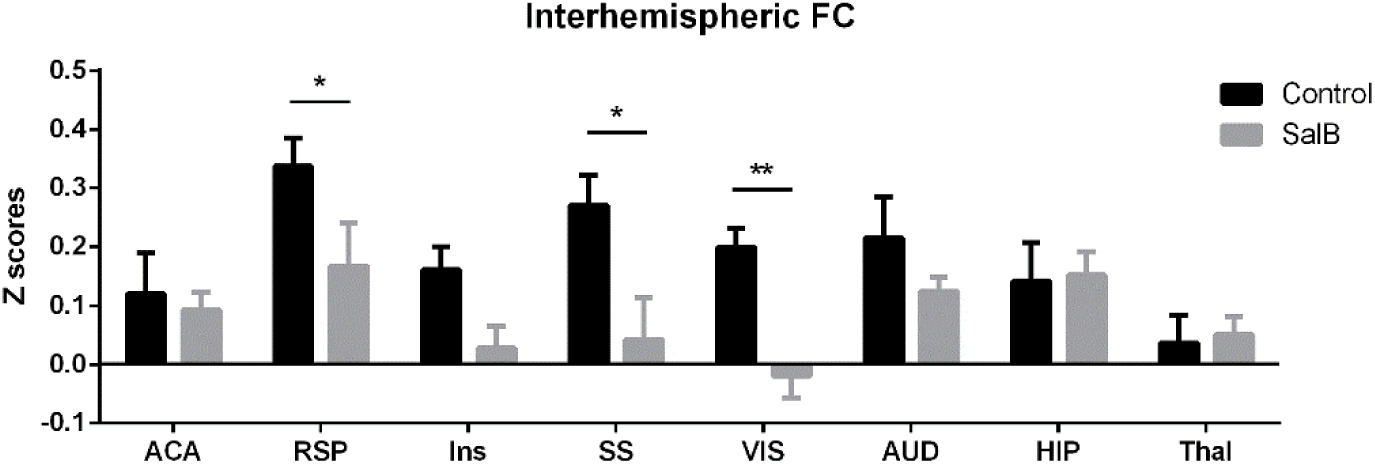
Inhibition of the right ACA induced decreased interhemispheric functional connectivity. Bar graphs show mean Z-scores ± SEM of all brain regions (anterior cingulate area (ACA), retrosplenial cortex (RSP), insular cortex (Ins), visual cortex (VIS), somatosensory cortex (SS), auditory cortex (AUD), hippocampus (HIP) and thalamus (Thal); Paired Student’s T-test). * p≤0.05, ** p≤0.01).

### 3.3 Physiological parameters

To exclude potential physiological effects of SalB and/or DMSO injections. The breathing rate was recorded during all *in vivo* imaging procedures. The breathing rates were compared across all groups to evaluate whether observed neuronal activity and FC changes are influenced by alterations of the animals’ physiological condition. As such, breathing rates were compared between the sham-treated group and the KORD-expressing group for the phMRI scans and between the two rsfMRI scan sessions, i.e. upon vehicle injection and SalB injection, in the KORD-expressing group. The breathing rate in the sham group (mean ± SEM: (155 ± 7) breaths per minute) did not significantly differ from the breathing rate of the KORD-expressing mice (mean ± SEM: (160 ± 4) breaths per minute; two-sample Student’s T-test, p=0.4). Furthermore, the injected SalB concentration did not significantly influence the breathing rate in the KORD – expressing mice (mean ± SEM (160 ± 4) breaths per minute with 3 mg/kg SalB injection and (165 ± 3) breaths per minute with 6 mg/kg SalB injection; Paired Student’s T-test p=0.4). In addition, no significant difference could be observed in the breathing rate between the two rsfMRI scan sessions in the KORD-expressing group (mean ± SEM: (157 ± 7) breaths per minute after vehicle injection and (158 ± 23) breaths per minute after SalB injection; Paired Student’s T-test, p=0.9).

## 4. Discussion

In this study, inhibitory chemogenetics were combined with non-invasive functional MRI to evaluate the effects of neuronal inhibition in the ACA on whole-brain neuronal activity and FC. Inhibitory KORD were expressed in the right ACA of mice, after which phMRI scans and rsfMRI scans were acquired. The phMRI results showed that inhibition of the neurons in the right ACA induced BOLD signal decreases as well as BOLD signal increases. Significantly decreased BOLD signals were observed in the target region, i.e. right ACA, as well as in the left ACA, right and left INS, left SS, right VIS and left AUD. On the other hand, increased BOLD signals were measured in the right SS, right and left AN, right RSP and right HIP. We further hypothesized that inhibition can induce desynchronization of neuronal firing which can lead to disrupted FC. Indeed, significantly decreased FC was observed for the ACA, RSP, SS, VIS, AUD and HIP. Histological examination of the brains, using uDISCO clearing and light-sheet microscopy, revealed KORD expression in the target region and structurally connected brain regions.

### 4.1 Neuronal inhibition of the right anterior cingulate area induced altered BOLD responses throughout the brain

The KORD-induced inhibition of the right ACA caused significant modulations in the BOLD signals in the ACA as well as inter-connected brain regions. A study has previously demonstrated that chemogenetic-induced neural activity modulations could be observed using phMRI in the mesocorticolimbic system [24]. To the best of our knowledge, the study presented here shows for the first time whole-brain network activity modulations upon the combined use of KORD-induced neuronal inhibition and phMRI in mice. This approach allowed us to gain a better understanding of the importance of a preserved neuronal activity in the ACA for whole-brain network activity and connectivity.

Although the exact function of the ACA is not yet fully known, several tracer injection studies have shown its involvement in various networks. The ACA has been shown to mediate the transfer of information between sensory areas, such as the somatosensory, visual and auditory cortices, and to higher order brain regions including the prefrontal cortex [34]. This suggests that the ACA is implicated in movement orientation and coordination of the eyes, head and body in object searching and spatial navigation. These connections of the ACA with sensory regions could also be observed in our KORD-phMRI results. As such, decreased BOLD signals were measured in the right visual cortex and left auditory cortex and increased BOLD signals were observed in the right somatosensory cortex. Furthermore, it has been shown that the ACA also receives information from the subiculum and transmits it to other prefrontal areas and it is suggested to be part of a network supporting spatial orientation and episodic memory [4, 34]. Furthermore, previous tracer injection studies have shown that the ACA has reciprocal connections with the retrosplenial cortex, visual cortex as well as with thalamic nuclei [35]. This is fully consistent with the observed BOLD signal alterations in these brain regions after inhibition of the ACA in our study. However, given that MRI is only an indirect measure of neuronal activity, and is unable to differentiate between excitatory and inhibitory cell types, future research will be necessary in order to clarify the contributions of specific cell classes in mediating the BOLD signal alterations.

### 4.2 Neuronal inhibition of the right anterior cingulate area induced whole-brain functional connectivity decreases

In this study, rsfMRI showed decreased FC in different networks during inactivation of the ACA. Previous human brain imaging studies used graph-theory based methods to identify a set of brain regions that play a central role in the functional topology of the human brain. These regions include cingulate cortex, insular cortex, frontal cortex, temporal cortex and the posterior cortex [36-38]. Similarly, Liska et al [39] found high strength nodes in the mouse brain including the cingulate cortex and the prefrontal cortex. This might explain the results in our study, as the inhibition of the ACA resulted in decreased FC in various brain regions. By evaluating the FC of both unilateral and contralateral seeds, we could assess whether the effect of KORD-mediated neuronal inhibition is localized in the ipsilateral, affected hemisphere or is also spreading to the contralateral side. Our findings demonstrated FC alterations in both the ipsilateral and contralateral hemisphere. We observed decreased FC for brain regions that are known to be structurally connected with the ACA (see above). This is in line with the emerging view that functional correlations in spontaneous brain activity are guided by underlying anatomical connections [14, 39]. The decreased FC can thus be due to neuronal firing desynchronization mediated by disturbed inputs coming from the ACA. We conjecture that the disruption of this hub region, with high connection strength and high degree of structural connectivity, leads to widespread alterations in the integrity of the correlations between brain networks and highlights the importance of hub node integrity in the dynamics of the whole network.

## 5. Conclusion

Our study demonstrates the ability of the combined use of KORD with *in vivo* MRI to assess large-scale network activity and FC in response to inhibition of a hub region, namely the right ACA in the mouse brain. We showed that KORD-induced inhibition of this area could induce BOLD signal decreases and increases in connected brain regions throughout both hemispheres. The affected brain regions are in line with previous tracer injection studies demonstrating projection areas of the ACA in mice. Furthermore, these connected brain regions showed decreased FC measures. To conclude, our study identifies the DREADD technology as a valuable and reliable tool that can be used in combination with non-invasive imaging methods. This approach might be of significant value for future neuroscientific research trying to unravel the underlying mechanisms of the function and dysfunction of neuronal networks.

