## Supplementary material for "Chemogenetic silencing of neurons in the mouse anterior cingulate area modulates neuronal activity and functional connectivity"

**Supplementary figure:**

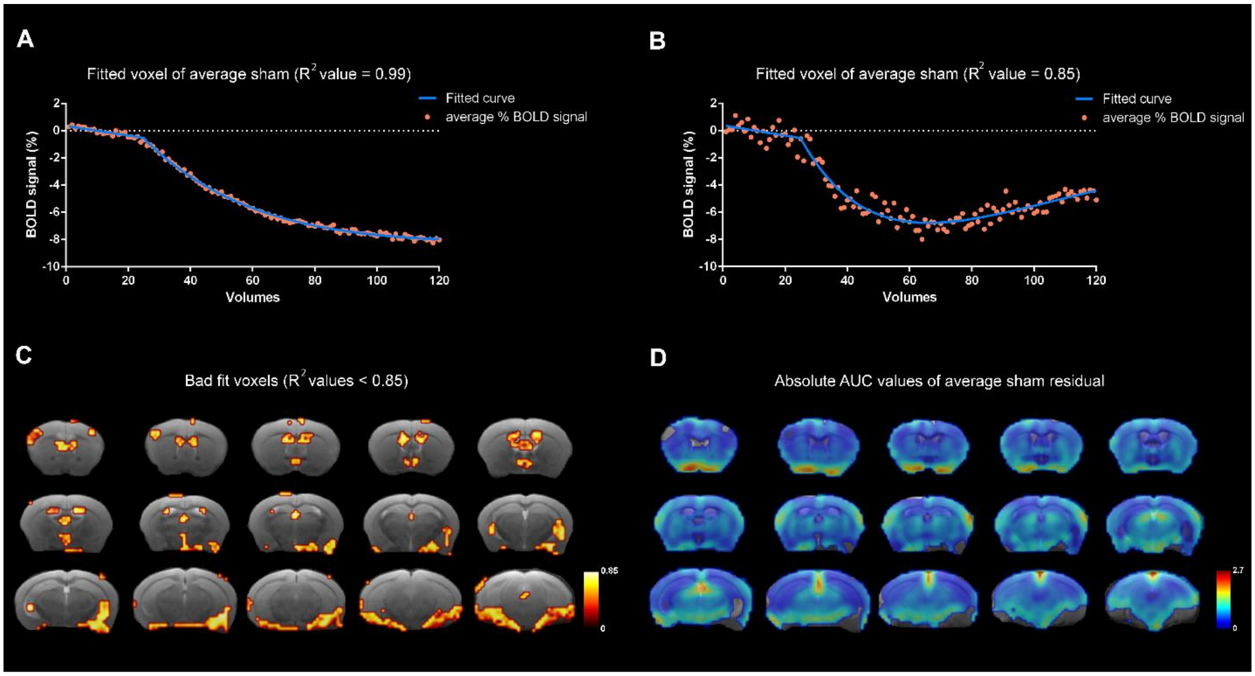


***Supplementary Figure 1. Estimating the non-specific effects of SalB and DMSO in the shams.*** *A + B) a linear polynomial fit and a double exponential fit were used to fit the average sham percent BOLD signal change. Graphs represent representative examples of a voxel with a good fit (A, $R^2$ value = 0.99) and a voxel with a less good fit (B, $R^2$ value = 0.85). All voxels with a $R^2$ value ≥ 0.85 are used for further analyses. The average sham percent BOLD signal change (blue) and the fitted curve (red) are shown C) Representation of the voxels with a bad fit ($R^2$ value < 0.85). These voxels are excluded from further analyses. D) Representation of the absolute AUC residuals of the shams after correction of the fitted curves.*

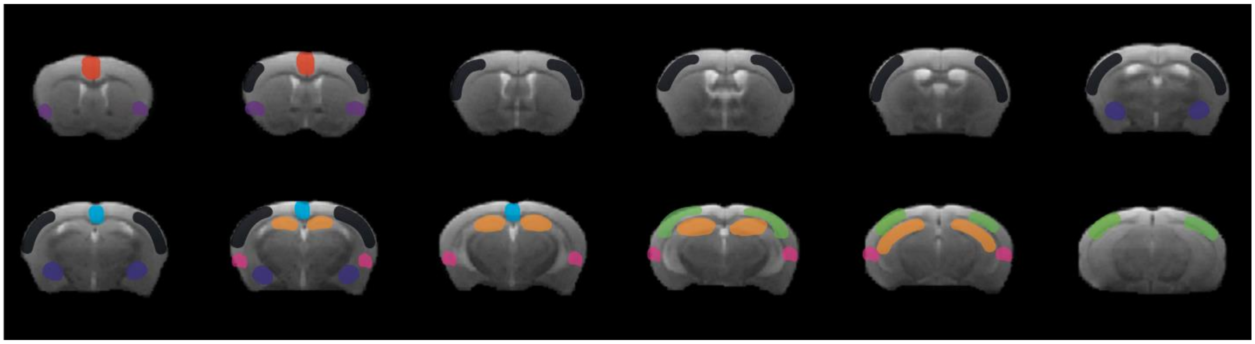

***Supplementary Figure 2. Illustration of the ROI selection for the phMRI analyses.*** *ROIs were selected for left and right hemisphere separately. Red: anterior cingulate area, purple: insular cortex, black: somatosensory cortex, magenta: auditory cortex, blue: amygdalar nuclei, orange: hippocampus, cyan: retrosplenial cortex, green: visual cortex.*

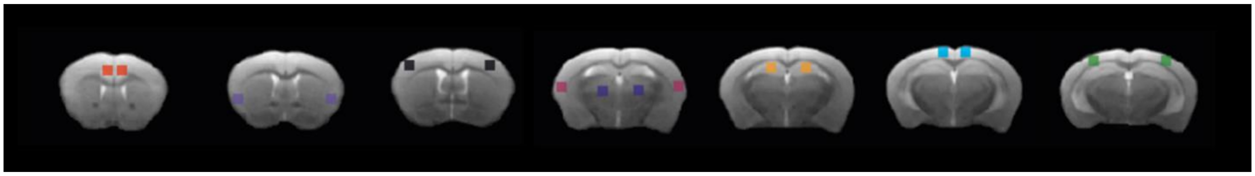

***Supplementary Figure 3. Illustration of the ROI selection for the rsfMRI analyses****. ROIs were selected for left and right hemisphere separately. Red: anterior cingulate area, purple: insular cortex, black: somatosensory cortex, magenta: auditory cortex, blue: thalamus, orange: hippocampus, cyan: retrosplenial cortex, green: visual cortex.*

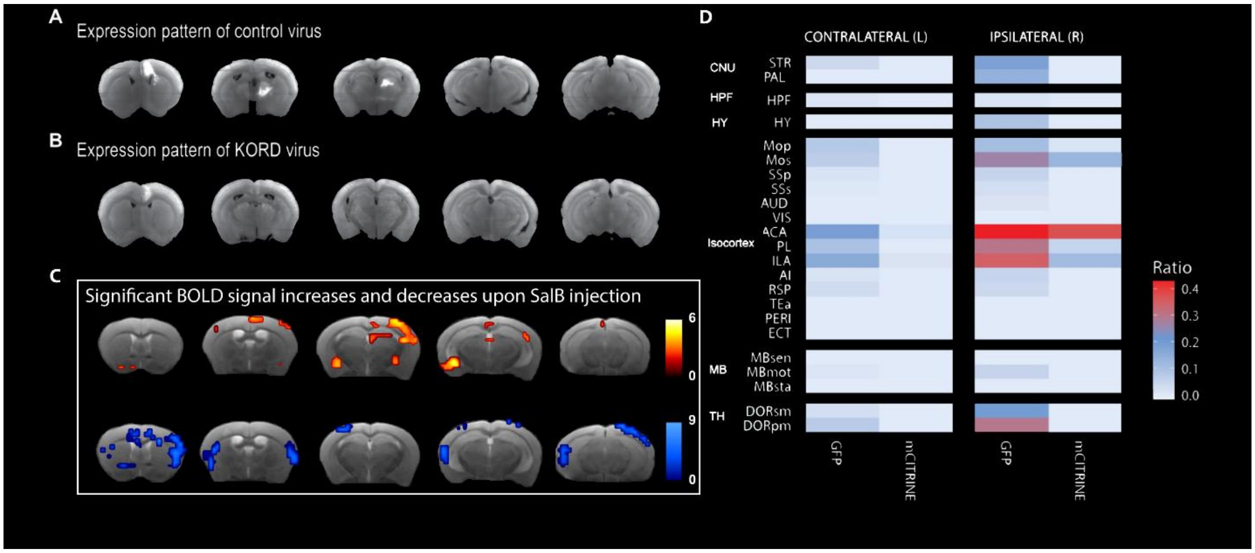


**Supplementary Figure 4.** *A) Representative example of the EGFP signal throughout the brain. B) Representative example of the spreading of the mCitrine signal throughout the brain. C) Illustration of the significant BOLD signal changes upon SalB injection measured by phMRI (same as Figure 1). D) Quantification of the mean EGFP and mCitrine signals in the brain. Color code represent the square root*

*of the ratio (%) of the positive voxels over the total amount of voxels in a ROI delineated based on the Allen brain Atlas. The targeted injection site overlaps with the ACA of the right hemisphere in both the sham group and the KORD expression group. Region based quantification of the mCitrine signal showed that the KORD expression was mainly localized to the target region (see also panel B for representative example). In contrast, the EGFP control virus could also be observed in neighboring regions, including motor and somatosensory regions, retrosplenial cortex, insular cortex, thalamic nuclei and caudate putamen, with spreading to the contralateral hemisphere. It's known that the mCitrine tends to produce a weaker signal compared to EGFP, which might explain the distinction between the spreading patterns of both fluorescent proteins. When comparing panels C and D, regions can be identified which show significant alterations in BOLD signal and EGFP/mCitrine expression. These regions include the ACA, somatosensory cortex, retrosplenial cortex and insular cortex.*

*Abbreviations: cerebral nuclei (CNU), striatum (STR), pallidum (PAL), hippocampal formation (HPF), hypothalamus (HY), primary motor area (MOp), secondary motor area (MOs), primary somatosensory area (SSp), supplemental somatosensory area (SSs), auditory areas (AUD), visual areas (VIS), anterior cingulate area (ACA), prelimbic area (PL), infralimbic area (ILA), agranular insular area (AI), retrosplenial area (RSP), temporal association areas (TEa), perirhinal area (PERI), ectorhinal area (ECT), midbrain (MD), midbrain sensory related (MBsen), midbrain motor related (MBmot), midbrain behavioral state related (MBsta), thalamus (TH), thalamus sensory-motor cortex related (DORsm), thalamus polymodal association cortex related (DORpm).*
